# The Prevalence and Patterns of Combined Psychotropic Polypharmacy among Acute Care Hospitals in Japan

**DOI:** 10.1101/649012

**Authors:** Sayuri Shimizu, Yasuyuki Okumura, Koichi B. Ishikawa, Shinya Matsuda, Hiroto Ito, Kiyohide Fushimi

## Abstract

**Purpose:** To evaluate the prescribing patterns of psychotropic polypharmacy for inpatients of acute care hospitals in Japan.

**Methods:** Administrative data on 2,639,885 patients admitted to acute care hospitals in Japan between July and December of 2008 were analyzed retrospectively. We defined psychotropic medications as antipsychotics, antidepressants, benzodiazepines, and other sedatives/hypnotics and studied their prescription patterns during the hospitalization of patients with stroke, acute cardiac infarction, cancer, and diabetes mellitus.

**Results:** At least one psychotropic drug was prescribed in 35.9% of all cases. Two-drug combinations of antipsychotic drugs were prescribed for stroke patients in 14,615 cases (1.4%), more than 2 in 3,132 cases (0.3%), and 22.4% of cases were prescribed 2 or more psychotropic drugs in addition to antipsychotic drugs. Amongst upper gastrointestinal cancer patients, 7.7% were prescribed a combination of 2 or more drugs, including benzodiazepines. Of the upper gastrointestinal cancer patients who were prescribed benzodiazepines, 20.3% were also prescribed 2 or more psychotropic drugs. Amongst stroke and upper gastrointestinal cancer patients, 36.6% and 35.6%, respectively, were treated with combination therapy using drugs of this class and others.

**Conclusion:** There is a pattern of polypharmacy that combines benzodiazepines and other sedatives/hypnotics with antidepressants or antipsychotic drugs, and this study provides a detailed analysis of this within acute care hospitals. Our results indicate the need for additional research into the efficacy of polypharmacy for inpatients in non-psychiatric settings.

## INTRODUCTION

Psychotropic polypharmacy constitutes a major burden on the health care system worldwide. Polypharmacy has several negative consequences associated with it, including the increased risk of medication related adverse events, higher mortality, poorer adherence, drug interactions, and greater costs.^1–5^ The current treatment guidelines strongly discourage psychotropic polypharmacy in routine clinical practice due to the lack of evidence regarding its efficacy and safety.^6–12^ Nevertheless, the concurrent use of multiple psychiatric drugs is a common practice, and such use has been increasing in several countries in psychiatric settings.^14–17^

The same tendency was found amongst inpatients in non-psychiatric settings. ^18–20^ Earlier studies suggested that polypharmacy was more common in elderly patients and inpatients with severe chronic diseases.^21, 22^ Often, inpatients receiving multiple medications for treatment of their conditions and psychological problems such as insomnia, anxiousness, and irritability are much less likely to receive psychosocial care. However, only a few studies have reported the prevalence of psychotropic polypharmacy in non-psychiatric inpatients.

There is much unknown about the comprehensive patterns of psychotropic polypharmacy. To date, most studies have addressed only same-class polypharmacy, ^23–26^ limited sets of drug polypharmacy, ^27^ and limited classes of polypharmacy.^28–30^ The lack of comprehensive studies on psychotropic polypharmacy patterns are partly due to the difficulty of analyzing data. In many clinical situations, a polypharmacy of different classes of psychotropic medications is indicated,^30^ and understanding these prescribing patterns should lead to improved patient safety, prescribing guidelines, and policymaking.

The present study was conducted to evaluate the prevalence and patterns of combination psychotropic polypharmacy among inpatients in a nationwide sample of acute care hospitals in Japan. To our knowledge, this is a first report to examine these prevalence and patterns in inpatients other than those on psychiatric hospitals.

## METHODS

### Data Source

Data were collected from the discharge claim records from Japanese acute care hospitals that were either utilizing or preparing to implement the Diagnosis Procedure Combination (DPC) code as a per-diem payment system in 2008.^31–33^ The data were voluntary offered to the DPC study group by 855 hospitals located throughout Japan that had agreed to participate. The 2008 version of the DPC/PDPS data included 18 major diagnostic categories and 506 disease subcategories coded in International Classification of Diseases and Related Health Problem, Tenth Revision (ICD-10). We used discharge data collected between July 1 and December 31, 2008. The study was approved by the Institutional Review Board of Tokyo Medical and Dental University.

### Participants

We extracted DPC/PDPS data for inpatients hospitalized for stroke, acute myocardial infarction (AMI), cancer, and diabetes mellitus (DM). Stroke, AMI, and cancer are the top 3 leading causes of mortality in Japan, accounting for approximately 60% of all deaths. For this study, we included only the more common types of cancer as they are associated with greater patient numbers, and these were cancers of the lung, upper and lower gastrointestinal tract, and breast (the most prevalent female cancer). DM was included because it has the greatest impact on overall health. The 4 diseases were defined as follows. Stroke (ICD-10 codes: I60, I61, I629, I63-66, I672, I675-682, I688, I690-694, I698, I978, G45-46, Q280-283), AMI (ICD-10 codes: I20-22, I24-25), DM (ICD-10 code: E10-14), lung cancer (ICD-10 codes: C33-34, C37, C381-383, C388, C39, C771, C780-781, C783, D021-022, D024), lower gastrointestinal cancer (ICD-10 codes: C17-21, C260, C268-269, C451, C480-482, C488, C772, C775, C784-785, D010-014, D017), gastric cancer (ICD-10 codes: C16, D002), breast cancer (ICD-10 codes: C50,D05).

### Psychotropic Medication Definition

We defined psychotropic medications in this study as antipsychotics, antidepressants, benzodiazepines, and other sedatives/hypnotics prescribed during hospitalization. We classified 92 drugs according to one of the most widely used prescription handbooks in the Japanese clinical setting. More information about the detailed classification of these drugs is provided in the supplemental table.

### Analytic Approach

Analyses were conducted in 2 stages, the first of which was devised to allow the handling of very large data sets containing millions of rows. The data examined in this study is a nationwide large-scale data set consisting of (1) a basic patient information file of 2,639,885 cases and (2) a psychotropic prescription file of 9,719,615 cases, to be analyzed in conjunction with anonymized patient IDs. SQL Server 2008 (Microsoft Corporation) software was therefore used to build up the database as a data-handling system. The second stage was devised to describe the demographic and patterns of concomitant use of psychotropic drugs from the same psychotropic medication class and of different classes. Our data included information on all psychiatric drugs covered by public health insurance, and therefore, this study did not involve sampling data but instead, studied an entire population. Therefore, it was unnecessary to extrapolate the results to all patients, and so, we used descriptive statistics. We calculated the proportion of patients with each disease type, who were prescribed one or more psychotropic drugs. Data were analyzed with the SQL Server 2008 and the JMP statistical package (9.0.2 Version).

## RESULTS

Figure 1 shows the fraction of psychotropic drug prescribed for each disease. At least one psychotropic drug was prescribed in 35.9% of those cases. Antipsychotics were prescribed in 235,540 cases (8.9%), antidepressants in 71,739 cases (2.7%), and benzodiazepines in 874,088 cases (33.1%). Other sedatives/hypnotics were prescribed in 852,809 cases (32.3%). Over 40% of all cancer cases were prescribed at least one psychotropic drug, 41.7% in lower gastrointestinal tract cancer, 47.7% in breast cancer, 48.0% in lung cancer, and 51.4% in upper gastrointestinal cancer. The prescription proportion of antipsychotic drugs was 3.3% in AMI. In contrast, the following cancers had a prescription proportion of more than 10% for antipsychotic drugs: lung cancer (14.1%), lower gastrointestinal tract cancer (10.7%), and upper gastrointestinal cancer (13.9%). The prescription of psychotropic drugs in diabetes was lower than that in other diseases, although antidepressants were prescribed in 3.0% of cases, which is the highest proportion of all 4 diseases. The prescriptions of other sedatives/hypnotics were high for upper gastrointestinal cancer (14.8%), lung cancer (18.1%), and breast cancer (19.6%).

**Figure 1.**
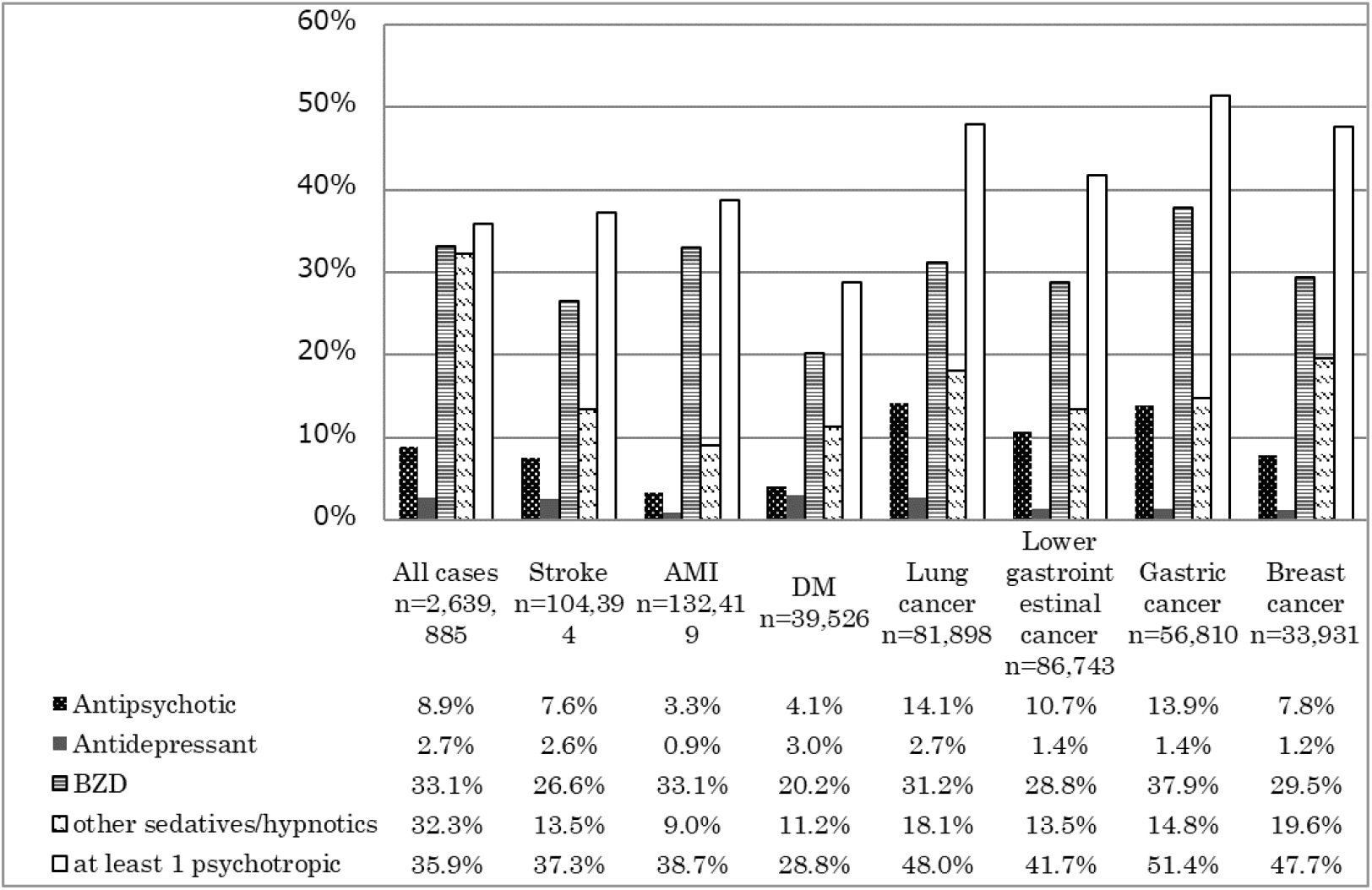
Prescription fraction of psychotropic drug by disease for inpatients at non-psychiatric acute care hospitals from July 2008 to December 2008. See supplemental section for description of medications in each class.

Table 1 shows the fractions of same-class psychotropic polypharmacy by disease. Regarding the polypharmacy of antipsychotic drugs, a 2-drug combination of antipsychotic drugs was prescribed for stroke patients in 14,615 cases (1.4%), and a combination of 3 or more antipsychotic drugs was prescribed in 3,132 cases (0.3%). Furthermore, 22.4% of the patients who were prescribed antipsychotic drugs were additionally prescribed 2 or more psychotropic drugs. Amongst cancer patients, 1,801 patients (2.2%) received a 2-drug combination, and 573 patients (0.7%) received a combination of 3 or more drugs. In addition, 20.6% of the patients who were prescribed antipsychotic drugs also received a combination of 2 or more psychotropic drugs. In diabetes, 0.5% of the patients received a combination of 2 or more antidepressants, accounting for 19.9 % of the cases in which antidepressants were prescribed. Amongst upper gastrointestinal cancer patients, 30.2% of the patients received a benzodiazepine as a single agent, and 7.7% of the patients received 2 or more drugs combined with benzodiazepines. Among the patients who were prescribed benzodiazepines, 20.3% additionally received 2 or more psychotropic drugs.

**Table1.** Analysis of same-class psychotropic polypharmacy prescribing by disease type for inpatients at non-psychiatric acute care hospitals between July and December 2008

|  | Antipsychotic |  |  | Antidepressant |  | BZD |  | other<br>sedatives/hypnotics |  |
| --- | --- | --- | --- | --- | --- | --- | --- | --- | --- |
|  | 1 | 2 | ≥3 | 1 | ≥2 | 1 | ≥2 | 1 | ≥2 |
| Stroke | 5.9% | 1.4% | 0.3% | 2.4% | 0.2% | 21.3% | 5.2% | 12.5% | 0.9% |
| AMI | 2.6% | 0.6% | 0.1% | 0.8% | 0.1% | 26.0% | 7.0% | 8.7% | 0.4% |
| DM | 3.3% | 0.6% | 0.3% | 2.5% | 0.5% | 15.1% | 5.1% | 10.8% | 0.5% |
| Cancer |  |  |  |  |  |  |  |  |  |
| Lung | 11.2% | 2.2% | 0.7% | 2.4% | 0.2% | 24.2% | 7.0% | 17.3% | 0.8% |
| Lower<br>gastrointestinal<br>tract | 9.1% | 1.2% | 0.3% | 1.3% | 0.2% | 23.3% | 5.5% | 13.0% | 0.5% |
| Gastric | 11.7% | 1.8% | 0.4% | 1.2% | 0.1% | 30.2% | 7.7% | 14.3% | 0.5% |
| Breast | 6.7% | 0.9% | 0.3% | 1.0% | 0.2% | 24.9% | 4.6% | 19.3% | 0.3% |

Antipsychotic drug prescriptions were classified into single-, double- and multiple-agent combinations. Antidepressants, benzodiazepine, and other sedatives/hypnotics were classified into groups of single- and multiple-agent prescription. See supplemental section for a detailed description of each class of medication.

Tables 2–3 list the 20 most prevalent patterns of prescribing psychotropic drug combinations for inpatients with stroke and upper gastrointestinal cancer, while those for other diseases are listed in the Supplemental Appendix. The 20 most frequently occurring patterns account for 94.3% of the total in stroke patients and 96.7% in patients with upper gastrointestinal cancer. In both cases, single-agent benzodiazepines or other sedatives/hypnotics, or a prescription regimen with a combination of these, were the top 4 most common prescribing patterns. It was also revealed that in cases of both upper gastrointestinal cancer and stroke, polypharmacy with drugs in the same class or in different classes are commonly used. Amongst stroke and upper gastrointestinal cancer patients, a psychotropic drug as a single agent was prescribed in 63.4 % and 64.4% of cases, respectively. The other prescriptions consisted of a combination therapy involving both drugs of the same class and of different classes. It is also noteworthy that there was a pattern of drug therapy, which combined benzodiazepines and other sedatives/hypnotics with antidepressants or antipsychotic drugs, and 5.8% of the stroke patients and 4.9% of the patients with upper gastrointestinal cancer received 3 or more different classes of polypharmacy.

**Table2.** The 20 most prevalent prescribing patterns of psychotropic drug combinations for stroke inpatients at non-psychiatric acute care hospitals between July and December 2008. (A total of 38,889 prescriptions amongst 104,394 cases)

| rank | n | % | Antipsychotic | Antidepressant | BZD | other<br>sedatives/hypnotics |
| --- | --- | --- | --- | --- | --- | --- |
| 1 | 15535 | 39.95% |  |  | 1 |  |
| 2 | 6627 | 17.04% |  |  |  | 1 |
| 3 | 2993 | 7.70% |  |  | 1 | 1 |
| 4 | 2479 | 6.37% |  |  | ≥2 |  |
| 5 | 1875 | 4.82% | 1 |  |  |  |
| 6 | 1481 | 3.81% | 1 |  | 1 |  |
| 7 | 813 | 2.09% |  |  | ≥2 | 1 |
| 8 | 660 | 1.70% |  | 1 |  |  |
| 9 | 629 | 1.62% | 1 |  |  | 1 |
| 10 | 618 | 1.59% | 1 |  | ≥2 |  |
| 11 | 506 | 1.30% | 1 |  | 1 | 1 |
| 12 | 436 | 1.12% |  | 1 | 1 |  |
| 13 | 305 | 0.78% | 2 |  | 1 |  |
| 14 | 303 | 0.78% |  |  |  | ≥2 |
| 15 | 295 | 0.76% | 1 |  | ≥2 | 1 |
| 16 | 283 | 0.73% | 2 |  |  |  |
| 17 | 258 | 0.66% |  |  | 1 | ≥2 |
| 18 | 208 | 0.53% |  | 1 | ≥2 |  |
| 19 | 199 | 0.51% | 2 |  | ≥2 |  |
| 20 | 185 | 0.48% |  | 1 |  | 1 |

See supplemental section for a detailed description of each class of medication.

**Table3.** The 20 most prevalent prescribing patterns of psychotropic drug combinations for inpatients with Gastric Cancer at non-psychiatric acute care hospitals between July and December 2008. (A total of 29,179 prescriptions amongst 56,810 cases)

| rank | n | % | Antipsychotic | Antidepressant | BZD | other<br>sedatives/hypnotics |
| --- | --- | --- | --- | --- | --- | --- |
| 1 | 12352 | 42.33% |  |  | 1 |  |
| 2 | 3948 | 13.53% |  |  |  | 1 |
| 3 | 2396 | 8.21% | 1 |  |  |  |
| 4 | 2167 | 7.43% |  |  | ≥2 |  |
| 5 | 1803 | 6.18% | 1 |  | 1 |  |
| 6 | 1765 | 6.05% |  |  | 1 | 1 |
| 7 | 742 | 2.54% | 1 |  | ≥2 |  |
| 8 | 637 | 2.18% | 1 |  |  | 1 |
| 9 | 510 | 1.75% |  |  | ≥2 | 1 |
| 10 | 470 | 1.61% | 1 |  | 1 | 1 |
| 11 | 271 | 0.93% | 1 |  | ≥2 | 1 |
| 12 | 239 | 0.82% | 2 |  | 1 |  |
| 13 | 225 | 0.77% | 2 |  |  |  |
| 14 | 180 | 0.62% | 2 |  | ≥2 |  |
| 15 | 96 | 0.33% |  | 1 |  |  |
| 16 | 96 | 0.33% |  | 1 | 1 |  |
| 17 | 94 | 0.32% |  |  |  | ≥2 |
| 18 | 86 | 0.29% | 2 |  | 1 | 1 |
| 19 | 76 | 0.26% | 2 |  |  | 1 |
| 20 | 74 | 0.25% | 2 |  | ≥2 | 1 |

See supplemental section for a detailed description of each class of medication.

## DISCUSSION

The key finding of this study is that psychotropic drugs are commonly prescribed to inpatients in non-psychiatric departments. Furthermore, combinations of these agents in the same class and polypharmacy with different classes have been prescribed to a broad range of patients. Some psychotropic drugs have been prescribed in all the medical institutions covered by this study. We therefore suggest that physicians in non-psychiatric departments should be given access to continuing education opportunities to extend their knowledge of psychotropic drug therapies.

On an average, 8.9% of all inpatients were prescribed antipsychotics, with the highest rate of prescription (14.1%) for lung cancer patients. Comparison of the fraction of prescription is difficult due to the lack of information on the prescription of psychotropic drugs to non-psychiatric inpatients. There is a possibility that some patients received antipsychotic drugs for various pathological conditions as an off-label use. Antipsychotic drugs have been used to deal with various mental symptoms such as anxiety, sleep disorders, and delirium, although there is only very limited evidence of their effectiveness, benefit, or safety for such uses. Therefore, further research is needed to evaluate that benefits and risks associated with the off-label use of antipsychotic drugs for non-psychiatry settings. It is also noteworthy that 20% of the cases were of antipsychotic polypharmacy prescription in a non-psychiatric department. There is also only limited evidence that polypharmacy of antipsychotic drugs is beneficial,^34^ and indeed, there is evidence for it exacerbating symptoms and causing adverse effects.^35–37^ Polypharmacy should be limited to patients with refractory schizophrenia, and it is recommended that only the minimum effective doses of antipsychotics should be prescribed. In this study, we also found a few cases of 3 or more antipsychotic drugs being prescribed in combination. These polypharmacy therapies should be carefully reconsidered.

The highest proportion of antidepressant prescription was 3.0% in diabetes patients, which is lower than that reported in previous studies. These earlier reports showed that the prevalence rate of comorbid major depressive disorder (based on diagnostic interviews) with diabetes was 11.4%, the presence of depressive symptoms (based on self-report scales) with diabetes was 31.0%,^38^ and depression with type 2 diabetes was 19.1%.^39^ Treatment with antidepressants requires long-term prescription and follow-up. Therefore, it might be difficult to start prescribing antidepressants for relatively short-term inpatient treatment in acute hospitals, and in addition, comorbid depression is often overlooked in non-psychiatric departments.^40, 41^ Nevertheless, hospitalization is also considered a unique opportunity to start treating depression that was previously untreated. A more vigorous approach to consultation-liaison psychiatry might encourage such action.

The proportion of patients prescribed sedatives/hypnotic is extremely high, with one-third of all inpatients having been administered these drugs. It was suggested that a number of specific prescribing patterns are chosen by clinicians for treating some inpatients with psychiatric symptoms with psychotropic drugs. These include benzodiazepine drugs as a base, followed by additional sedatives/hypnotics and then other antidepressants or antipsychotic drugs. The majority of the sedatives/hypnotics combine with benzodiazepine or the binding site of barbital and therefore have a similar mechanism of action. Thus, a combination of these drugs can be considered at a practical level to be a mass prescription. These drugs have a wide range of safe dosages, and severe side effects are rare. However, it should be noted that if these drugs are additionally prescribed aimlessly based on patients’ complaints, it could easily lead to mass multi-drug prescription. In addition, amongst the prescribed psychotropic drug combinations for all inpatients (appendix A3), benzodiazepines were the most frequently included. Since many prescriptions of etizolam and triazolam are unique to Japan, there are few previous studies that we can refer to for long-term safety data. A relatively high proportion of benzodiazepine prescriptions in Japan has also been previously reported.^42^ Therefore, it may be necessary to consider whether the prescription is beneficial, rather than focusing on the side effects such as dependence and withdrawal symptoms.

Our study has several limitations. When investigating the prescription of psychotropic drugs during a hospital stay, cases where the drug was switched to another in the same class were classified as a 2-drug combination case. Thus, we have conducted an additional analysis to compare the first prescription with the last one. The result of this indicated that there was only a low probability of a significant drug change. Further potential problems are that the findings are limited to inpatient admissions to acute care hospitals and the proportion of prescriptions in this study may have been underestimated. The latter may be because the acute hospitals that were surveyed have adopted a per-diem payment system for the hospitalization medical cost. Therefore, it is possible that patients might have bought the drugs that were prescribed and taken them before hospitalization to reduce costs.

In this study, “real world” comprehensive prescription patterns of psychotropic drugs for inpatients with stroke, AMI, DM, and 4 types of cancer were identified. There is a pattern of polypharmacy that combines benzodiazepines and other sedatives/hypnotics with antidepressants or antipsychotic drugs, and this study provides a detailed analysis of this within acute care hospitals. Our results indicate the need for additional research into the efficacy of polypharmacy for inpatients in non-psychiatric settings.

## ACKNOWLEDGEMENTS

Funding for the administration of this study was provided by Grants-in-Aid for Research on Policy Planning and Evaluation (Japanese Ministry of Health, Labour and Welfare H22-SEISAKU-SITEI-031 and H24-SEISAKU-SITEI-012). Financial support for writing this article was provided by a Health Labour Sciences Research Grant (H22-IYAKU-IPPAN-013) from the Ministry of Health, Labour and Welfare of Japan.

## Supplemental Digital Contents

### Supplemental Appendix A1. Psychotropic drug definition

We defined the use of psychotropic medications as the prescription of at least one substance among antipsychotics, antidepressants, benzodiazepines, and other sedatives/hypnotics. The details are as follows:

| Type | Generic name | Number of drugs |
| --- | --- | --- |
| antidepressants | Amitriptyline | 23 |
|  | Amoxapine | 4 |
|  | Clomipramine | 4 |
|  | Dosulepin | 2 |
|  | Fluvoxamine | 6 |
|  | Imipramine | 8 |
|  | Lofepramine | 2 |
|  | Maprotiline | 11 |
|  | Mianserin | 2 |
|  | Milnacipran | 24 |
|  | Nortriptyline | 3 |
|  | Paroxetine | 2 |
|  | Sertraline | 2 |
|  | Setiptiline | 4 |
|  | Trazodone | 6 |
|  | Trimipramine | 3 |
| Subtotal |  | 106 |
| antipsychotics | Aripiprazole | 4 |
|  | Blonanserin | 3 |
|  | Bromperidol | 24 |
|  | Carpipramine | 6 |
|  | Chlorpromazine | 32 |
|  | chlorpromazine-promethazine-combined | 2 |
|  | Clocapramine | 16 |
|  | Clothiapine | 3 |
|  | Fluphenazine | 10 |
|  | Haloperidol | 87 |
|  | haloperidol decanoate | 4 |
|  | Levomepromazine | 34 |
|  | Moperone | 2 |
|  | Mosapramine | 5 |
|  | Nemonapride | 3 |
|  | Olanzapine | 6 |
|  | Oxypertine | 4 |
|  | Perazine | 3 |
|  | Perospirone | 5 |
|  | Perphenazine | 29 |
|  | Pimozide | 4 |
|  | Pipamperone | 4 |
|  | Prochlorperazine | 7 |
|  | Propericiazine | 16 |
|  | Quetiapine | 3 |
|  | Risperidone | 78 |
|  | Spiperone | 5 |
|  | Sulpiride | 106 |
|  | Sultopride | 12 |
|  | Thioridazine | 5 |
|  | Tiapride | 36 |
|  | Timiperone | 11 |
|  | Trifluoperazine | 4 |
|  | Zotepine | 15 |
| Subtotal |  | 588 |
| benzodiazepines | Alprazolam | 10 |
|  | Bromazepam | 11 |
|  | Brotizolam | 27 |
|  | Chlordiazepoxide | 18 |
|  | clorazepate dipotassium | 1 |
|  | Clotiazepam | 15 |
|  | Cloxazolam | 3 |
|  | Diazepam | 68 |
|  | Estazolam | 6 |
|  | ethyl loflazepate | 18 |
|  | Etizolam | 39 |
|  | Fludiazepam | 4 |
|  | Flunitrazepam | 13 |
|  | Flurazepam | 7 |
|  | Flutazolam | 4 |
|  | Flutoprazepam | 3 |
|  | Haloxazolam | 4 |
|  | Lorazepam | 15 |
|  | Lormetazepam | 2 |
|  | Medazepam | 14 |
|  | Mexazolam | 4 |
|  | Nimetazepam | 4 |
|  | Nitrazepam | 27 |
|  | Oxazolam | 14 |
|  | Prazepam | 3 |
|  | Quazepam | 14 |
|  | Rilmazafone | 4 |
|  | Triazolam | 17 |
|  | Subtotal | 369 |
| other<br>sedatives/hypnotics | Amobarbital | 7 |
|  | Barbital | 7 |
|  | Bromovalerylurea | 17 |
|  | calcium bromide | 8 |
|  | Chloral | 5 |
|  | Passiflamin | 1 |
|  | pentobarbital calcium | 2 |
|  | Phenobarbital | 37 |
|  | phenobarbital sodium | 7 |
|  | secobarbital sodium | 2 |
|  | tandospirone citrate | 13 |
|  | Triclofos | 2 |
|  | Zolpidem | 2 |
|  | Zopiclone | 14 |
|  | Subtotal | 124 |

### Supplemental Appendix A2

The 20 most prevalent prescribing patterns of psychotropic drug combinations for AMI, DM, lung cancer, lower gastrointestinal cancer, and breast cancer patients.

**S1.** The 20 most prevalent prescribing patterns of psychotropic drug combinations for inpatients with AMI at non-psychiatric acute care hospitals between July and December 2008. (A total of 51,216 prescriptions amongst 132,419 cases).

| rank | n | % | Antipsychotics | Antidepressants | BZD | other sedatives/hypnotics |
| --- | --- | --- | --- | --- | --- | --- |
| 1 | 28910 | 56.4% |  |  | 1 |  |
| 2 | 6719 | 13.1% | | | $\geq 2$ | |
| 3 | 5806 | 11.3% |  |  |  | 1 |
| 4 | 3324 | 6.5% |  |  | 1 | 1 |
| 5 | 968 | 1.9% | 1 |  | 1 |  |
| 6 | 892 | 1.7% | | | $\geq 2$ | 1 |
| 7 | 717 | 1.4% | 1 |  |  |  |
| 8 | 645 | 1.3% | 1 | | $\geq 2$ | |
| 9 | 328 | 0.6% | 1 |  | 1 | 1 |
| 10 | 309 | 0.6% | 1 |  |  | 1 |
| 11 | 237 | 0.5% |  | 1 | 1 |  |
| 12 | 220 | 0.4% | 1 | | $\geq 2$ | 1 |
| 13 | 210 | 0.4% | | 1 | $\geq 2$ | |
| 14 | 186 | 0.4% | | | 1 | $\geq 2$ |
| 15 | 177 | 0.3% | 2 |  | 1 |  |
| 16 | 158 | 0.3% |  | 1 |  |  |
| 17 | 144 | 0.3% | 2 | | $\geq 2$ | |
| 18 | 117 | 0.2% | | | | $\geq 2$ |
| 19 | 97 | 0.2% | 2 |  |  |  |
| 20 | 85 | 0.2% | 2 | | $\geq 2$ | 1 |

See supplemental section for a detailed description of each class of medication.

**S2.** The 20 most prevalent prescribing patterns of psychotropic drug combinations for inpatients with DM at non-psychiatric acute care hospitals between July and December 2008. (A total of 11,386 prescriptions amongst 39,526 cases.)

| rank | N | % | Antipsychotic | Antidepressant | BZD | other<br>sedatives/hypnotics |
| --- | --- | --- | --- | --- | --- | --- |
| 1 | 4322 | 37.96% |  |  | 1 |  |
| 2 | 2451 | 21.53% |  |  |  | 1 |
| 3 | 923 | 8.11% | | | $\geq 2$ | |
| 4 | 858 | 7.54% |  |  | 1 | 1 |
| 5 | 355 | 3.12% | 1 |  |  |  |
| 6 | 284 | 2.49% | | | $\geq 2$ | 1 |
| 7 | 251 | 2.20% | 1 |  | 1 |  |
| 8 | 229 | 2.01% |  | 1 |  |  |
| 9 | 169 | 1.48% |  | 1 | 1 |  |
| 10 | 149 | 1.31% | 1 | | $\geq 2$ | |
| 11 | 115 | 1.01% | | 1 | $\geq 2$ | |
| 12 | 114 | 1.00% | 1 |  |  | 1 |
| 13 | 72 | 0.63% | 1 |  | 1 | 1 |
| 14 | 65 | 0.57% | 1 | 1 | 1 |  |
| 15 | 61 | 0.54% |  | 1 |  | 1 |
| 16 | 57 | 0.50% | | | | $\geq 2$ |
| 17 | 57 | 0.50% | | 1 | $\geq 2$ | 1 |
| 18 | 57 | 0.50% |  | 1 | 1 | 1 |
| 19 | 54 | 0.47% | 1 | | $\geq 2$ | 1 |
| 20 | 40 | 0.35% | 2 | | $\geq 2$ | |

See supplemental section for a detailed description of each class of medication.

**S3.** The 20 most prevalent prescribing patterns of psychotropic drug combinations for inpatients with lung cancer at non-psychiatric acute care hospitals between July and December 2008. (A total of 39,298 prescriptions amongst 81,898 cases.)

| rank | n | % | Antipsychotic | Antidepressant | BZD | other<br>sedatives/hypnotics |
| --- | --- | --- | --- | --- | --- | --- |
| 1 | 13025 | 33.14% |  |  | 1 |  |
| 2 | 7619 | 19.39% |  |  |  | 1 |
| 3 | 3479 | 8.85% | 1 |  |  |  |
| 4 | 2483 | 6.32% | | | $\geq 2$ | |
| 5 | 2433 | 6.19% |  |  | 1 | 1 |
| 6 | 2066 | 5.26% | 1 |  | 1 |  |
| 7 | 1054 | 2.68% | 1 |  |  | 1 |
| 8 | 747 | 1.90% | 1 | | $\geq 2$ | |
| 9 | 722 | 1.84% | | | $\geq 2$ | 1 |
| 10 | 638 | 1.62% | 1 |  | 1 | 1 |
| 11 | 414 | 1.05% | 2 |  |  |  |
| 12 | 391 | 0.99% | 2 |  | 1 |  |
| 13 | 348 | 0.89% | 1 | | $\geq 2$ | 1 |
| 14 | 268 | 0.68% |  | 1 |  |  |
| 15 | 263 | 0.67% | 2 | | $\geq 2$ | |
| 16 | 239 | 0.61% |  | 1 | 1 |  |
| 17 | 171 | 0.44% | | 1 | $\geq 2$ | |
| 18 | 169 | 0.43% | | | | $\geq 2$ |
| 19 | 158 | 0.40% | 2 |  | 1 | 1 |
| 20 | 144 | 0.37% | 2 |  |  | 1 |

See supplemental section for a detailed description of each class of medication.

**S4.** The 20 most prevalent prescribing patterns of psychotropic drug combinations for inpatients with lower gastrointestinal cancer at non-psychiatric acute care hospitals between July and December 2008. (A total of 29,200 prescriptions amongst 56,810 cases).

| rank | n | % | Antipsychotic | Antidepressant | BZD | other<br>sedatives/hypnotics |
| --- | --- | --- | --- | --- | --- | --- |
| 1 | 14748 | 37.96% |  |  | 1 |  |
| 2 | 6295 | 21.53% |  |  |  | 1 |
| 3 | 3164 | 8.11% | 1 |  |  |  |
| 4 | 2305 | 7.54% | | | $\geq 2$ | |
| 5 | 2037 | 3.12% |  |  | 1 | 1 |
| 6 | 1890 | 2.49% | 1 |  | 1 |  |
| 7 | 822 | 2.20% | 1 |  |  | 1 |
| 8 | 699 | 2.01% | 1 | | $\geq 2$ | |
| 9 | 643 | 1.48% | | | $\geq 2$ | 1 |
| 10 | 536 | 1.31% | 1 |  | 1 | 1 |
| 11 | 298 | 1.01% | 1 | | $\geq 2$ | 1 |
| 12 | 230 | 1.00% | 2 |  |  |  |
| 13 | 230 | 0.63% | 1 |  | 1 |  |
| 14 | 204 | 0.57% |  | 1 |  |  |
| 15 | 199 | 0.54% |  | 1 | 1 |  |
| 16 | 199 | 0.50% | 2 | | $\geq 2$ | |
| 17 | 107 | 0.50% | | | | $\geq 2$ |
| 18 | 96 | 0.50% | 2 |  | 1 | 1 |
| 19 | 93 | 0.47% | | 1 | $\geq 2$ | |
| 20 | 87 | 0.35% | 1 | 1 | 1 |  |

See supplemental section for a detailed description of each class of medication.

**S5.** The 20 most prevalent prescribing patterns of psychotropic drug combinations for inpatients with breast cancer at non-psychiatric acute care hospitals between July and December 2008. (A total of 16,185 prescriptions amongst 33,931 cases.)

| rank | n | % | Antipsychotic | Antidepressant | BZD | other<br>sedatives/hypnotics |
| --- | --- | --- | --- | --- | --- | --- |
| 1 | 6559 | 40.50% |  |  | 1 |  |
| 2 | 4597 | 28.38% |  |  |  | 1 |
| 3 | 1037 | 6.40% | 1 |  |  |  |
| 4 | 1008 | 6.22% |  |  | 1 | 1 |
| 5 | 820 | 5.06% | | | $\geq 2$ | |
| 6 | 493 | 3.04% | 1 |  | 1 |  |
| 7 | 269 | 1.66% | 1 |  |  | 1 |
| 8 | 228 | 1.41% | | | $\geq 2$ | 1 |
| 9 | 150 | 0.93% | 1 | | $\geq 2$ | |
| 10 | 135 | 0.83% | 1 |  | 1 | 1 |
| 11 | 77 | 0.48% | $\geq 2$ | | | |
| 12 | 71 | 0.44% | | | | $\geq 2$ |
| 13 | 60 | 0.37% | | 1 | $\geq 2$ | |
| 14 | 57 | 0.35% | $\geq 2$ | | 1 | |
| 15 | 49 | 0.30% | 1 | | $\geq 2$ | 1 |
| 16 | 45 | 0.28% |  | 1 |  |  |
| 17 | 44 | 0.27% | $\geq 2$ | | $\geq 2$ | |
| 18 | 41 | 0.25% |  | 1 | 1 |  |
| 19 | 32 | 0.20% | $\geq 2$ | | $\geq 2$ | 1 |
| 20 | 30 | 0.19% | $\geq 2$ | | | 1 |

See supplemental section for a detailed description of each class of medication.

### Supplemental Appendix A3

**S6.** The 50 most prescribed psychotropic drug combinations for all inpatients at non-psychiatric acute care hospitals between July and December 2008. (Total number of prescriptions = 947,006)

| Rank | N of hospitals | N of cases | % | N of Agents | Prescription combinations |
| --- | --- | --- | --- | --- | --- |
| 1 | 854 | 124,797 | 13.2% | 1 | Diazepam |
| 2 | 795 | 121,601 | 12.8% | 1 | Brotizolam |
| 3 | 826 | 107,637 | 11.4% | 1 | Zolpidem |
| 4 | 693 | 49,864 | 5.3% | 1 | Zopiclone |
| 5 | 488 | 42,008 | 4.4% | 1 | Rilmazafone |
| 6 | 845 | 29,851 | 3.2% | 1 | Etizolam |
| 7 | 686 | 23,326 | 2.5% | 1 | Flunitrazepam |
| 8 | 687 | 19,888 | 2.1% | 1 | Triazolam |
| 9 | 818 | 18,002 | 1.9% | 1 | Haloperidol |
| 10 | 695 | 12,292 | 1.3% | 2 | diazepam+brotizolam |
| 11 | 651 | 12,183 | 1.3% | 1 | Prochlorperazine |
| 12 | 456 | 10,386 | 1.1% | 1 | Triclofos |
| 13 | 711 | 9,972 | 1.1% | 2 | brotizolam+zolpidem |
| 14 | 684 | 9,137 | 1.0% | 2 | diazepam+zolpidem |
| 15 | 719 | 6,706 | 0.7% | 2 | etizolam+brotizolam |
| 16 | 669 | 6,522 | 0.7% | 1 | Phenobarbital |
| 17 | 569 | 6,441 | 0.7% | 1 | nitrazepam |
| 18 | 659 | 6,103 | 0.6% | 2 | haloperidol+brotizolam |
| 19 | 556 | 5,940 | 0.6% | 1 | estazolam |
| 20 | 740 | 5,933 | 0.6% | 2 | etizolam+zolpidem |
| 21 | 655 | 5,086 | 0.5% | 1 | risperidone |
| 22 | 614 | 4,574 | 0.5% | 1 | clotiazepam |
| 23 | 681 | 4,464 | 0.5% | 1 | alprazolam |
| 24 | 525 | 3,954 | 0.4% | 2 | brotizolam+zopiclone |
| 25 | 619 | 3,904 | 0.4% | 2 | haloperidol+zolpidem |
| 26 | 475 | 3,551 | 0.4% | 2 | diazepam+zopiclone |
| 27 | 493 | 3,460 | 0.4% | 2 | brotizolam+flunitrazepam |
| 28 | 733 | 3,438 | 0.4% | 1 | sulpiride |
| 29 | 692 | 3,352 | 0.4% | 2 | haloperidol+diazepam |
| 30 | 222 | 3,332 | 0.4% | 1 | lormetazepam |
| 31 | 465 | 3,315 | 0.4% | 2 | flunitrazepam+zolpidem |
| 32 | 547 | 3,216 | 0.3% | 1 | ethyl loflazepate |
| 33 | 289 | 3,124 | 0.3% | 2 | diazepam+rilmazafone |
| 34 | 631 | 3,086 | 0.3% | 2 | diazepam+etizolam |
| 35 | 514 | 3,049 | 0.3% | 2 | zolpidem+zopiclone |
| 36 | 563 | 2,939 | 0.3% | 2 | diazepam+phenobarbital |
| 37 | 691 | 2,859 | 0.3% | 1 | paroxetine |
| 38 | 440 | 2,513 | 0.3% | 2 | diazepam+triazolam |
| 39 | 331 | 2,498 | 0.3% | 2 | rilmazafone+zolpidem |
| 40 | 441 | 2,434 | 0.3% | 2 | haloperidol+flunitrazepam |
| 41 | 376 | 2,433 | 0.3% | 1 | lorazepam |
| 42 | 511 | 2,289 | 0.2% | 2 | etizolam+zopiclone |
| 43 | 425 | 2,138 | 0.2% | 2 | prochlorperazine+brotizolam |
| 45 | 315 | 2,084 | 0.2% | 2 | brotizolam+rilmazafone |
| 44 | 544 | 2,084 | 0.2% | 1 | amitriptyline |
| 46 | 419 | 2,083 | 0.2% | 2 | prochlorperazine+zolpidem |
| 47 | 447 | 2,012 | 0.2% | 1 | chlorpromazine |
| 48 | 438 | 1,989 | 0.2% | 2 | haloperidol+zopiclone |
| 49 | 269 | 1,944 | 0.2% | 2 | chloral+triclofos |
| 50 | 45 | 1,943 | 0.2% | 2 | diazepam+pentobarbital calcium |

